# Adverse Social Experiences in Adolescent Rats Results in Persisting Sex-Dependent Effects on Alcohol-Seeking Behavior

**DOI:** 10.1101/2020.12.07.411306

**Authors:** Akseli Surakka, Valentina Vengeliene, Ivan Skorodumov, Marcus Meinhardt, Anita C. Hansson, Rainer Spanagel

## Abstract

**Background:** Accumulating clinical evidence suggests women with prior exposure to adverse childhood experiences are more susceptible to alcohol relapse and other health-related issues. Yet, preclinical studies investigating sex-dependent effects of adolescent adverse social experiences (ASEs) on later alcohol-seeking behavior are lacking. This is mainly due to a lack of valid animal models and a shortage of studies comparing sexes. Therefore, we sought to investigate the sex-dependent effects of ASE on adult alcohol-seeking behavior, locomotion and reward sensitivity in both male and female rats.

**Methods:** We recently developed a rat model for adolescent peer-rejection which allows us to study the long-term consequences of ASEs. Peer-rejection interferes with adolescent rats‘ability to engage in adequate and reciprocal play behaviors that result in persistent dysregulation of social and pain-related behavior. Adolescent Wistar rats were reared from postnatal day (pd) 21 to pd 50 either within a group of Fischer 344 rats (ASE) or with Wistar rats (control). Adult male and female rats were tested in the reinstatement paradigm for cue-induced alcohol-seeking behavior, circadian locomotor activity, and sucrose consumption in adulthood long-after the termination of the peer-rejection condition.

**Results:** Peer-rejection induced persistent sex-dependent changes to cue-induced reinstatement. Females showed an increased reinstatement effect while peer-rejected males demonstrated a decrease. No differences were observed in circadian locomotor activity or reward sensitivity to sucrose.

**Conclusions:** Peer-rejection has lasting sex-dependent consequences on alcohol-seeking behavior without affecting locomotion or sweet reward sensitivity. Our results suggest that peer-rejected female rats represent a vulnerable population to study relapse-like behaviors similar to clinical findings. While males seem to buffer the peer-rejection effect and demonstrate resilience to later-life alcohol-seeking behaviors, measured by the reinstatement effect. Finally, we provide a novel approach to investigate the molecular and neurobiological underpinnings of ASEs on alcohol and other drug-seeking behaviors.

## Introduction

Increasing evidence from human studies show that adverse childhood experiences (ACEs) can result in altered brain structure and function (reviewed in Herzog and Schmahl, 2018). These changes, in turn, may result in several mental health issues later in life. The impact of multiple ACEs is best illustrated by the fact that the risk for suicide attempts is massively increased with an odds ratio of almost 18 (Hughes et al., 2019). ACEs can also alter reward processing (Hanson et al., 2016) and cause sleep disturbances (Hasler et al., 2012) that in turn can lead to alcohol-related problems such as early alcohol drinking initiation, alcohol binging, and problem alcohol use (lifetime) (Bellis et al., 2019; Benjet et al., 2016; Erdozain et al., 2015; Hughes et al., 2019, 2017). A large body of evidence has reported that women and men respond differently to ACEs and alcohol-related problems (Craig et al., 2019; Cunradi et al., 2020; Edwards et al., 2003; Fang and McNeil, 2017; Peltier et al., 2019). Specifically, women may have a larger risk for relapse to alcohol (Heffner et al., 2011; Hyman et al., 2008; Kennedy et al., 2013). In experimental settings women also demonstrate a greater cue reactivity to alcohol (Hartwell and Ray, 2013). Together these findings suggest that ACEs alter brain structure and function in a manner that increases susceptibility to alcohol-related problems and there seems to be gender-specific effects. It is thus of high relevance to develop a translational approach that allows to study the underlying molecular and neurobiological mechanisms that mediate the effects of ACEs on alcohol-related problems later in life. Here we set out to study this relationship in an appropriate animal model where male and female rats undergo adverse social experiences (ASEs) across adolescence and are then exposed to voluntary alcohol-self-administration and cue-induced reinstatement behavior in adulthood.

Theoretical models have proposed that individuals with ACEs are at an increased risk of later alcohol-related problems due to latent vulnerability (McCrory and Mayes, 2015). The core idea posits that the response to adversity although initially adaptive become maladaptive in the long-term. For example, being rejected by your peers during adolescence is distressful and may initially lead to devaluing social-contact to protect ones self-worth. In the long-term, peer-rejection may lead to less social contact and therefore less adaptive social skills later in life that could in turn increase the risk of later alcohol-related problems. Indeed, research has indicated that peer-rejection during adolescence is associated with future somatic, mental health issues including substance use (Finkelhor et al., 2015; Pinchevsky et al., 2014). Potential contributors to this increased risk of later alcohol-related problems are ACE induced; circadian dysregulation and reward system alterations (Hanson et al., 2016; Hasler et al., 2012). Although emerging evidence supports a relationship, there is still a significant gap in our understanding of the mechanisms to explain the association between peer-rejection and later alcohol-related problems. To explore the relationship between peer-rejection and later alcohol-related problems, we used our clinically relevant rat model of adolescent peer-rejection (Schneider et al., 2014, 2016a). Our peer-rejection-like ASE model moderates adolescent rat’s ability to properly engage in adequate and reciprocal play. Compared to other ASE models, that often use total social deprivation, our model allows animals to interact at their choice, better mirroring aspects of human peer-rejection. We have previously reported a variety of persistent behavioral (social interactions, social memory, pain-sensitivity and processing of socially transmitted information) and neurochemical (endocannabinoid system) alterations in peer-rejected rats (Schneider et al., 2014, 2016a). Considering that endocannabinoid alterations are common in individuals with alcohol-problems (Spanagel, 2020), it is plausible that peer-rejected rats, that also show a similar change in the endocannabinoid system, may demonstrate changes in their alcohol-seeking behaviors.

The goal of the present study was to determine whether peer-rejection-like ASE induces changes to alcohol self-administration and reinstatement of alcohol-seeking behavior in a sex-specific manner. Here we subjected male and female rats to peer-rejection across adolescence (PD 21 – 50) and then exposed them to voluntary alcohol self-administration in adulthood. We tested both male and female rats in the cue-induced reinstatement test to assess for relapse-like behavior. As mentioned earlier, circadian dysregulation is directionally associated with ACEs, future poor mental health (Walker et al., 2020), and alcohol use (Hasler et al., 2012). As our model recapitulates several of these aspects (ACEs and alcohol use), we evaluated whether peer-rejected rats would show circadian dysregulation assessed by the home-cage circadian locomotor activity. Finally, peer-rejection has been strongly associated with the development of depression of which anhedonia is a common feature (Slavich et al., 2010). We used the sucrose consumption test to assess anhedonia-like behavior in peer-rejected rats.

## Materials and Methods

### Animals

Forty 21-days-old female Wistar rats (Envigo, Venray Netherlands), twenty-four, 21-days-old male Wistar rats (Envigo, Venray, Netherlands), one-hundred and twenty, 21-days-old female Fischer 344 rats (Charles River, Sulzfeld, Germany) and seventy-two 21-days-old male Fischer 344 (Charles River, Sulzfeld, Germany) were used. All animals were housed in groups of four throughout their adolescence into adulthood (unless stated otherwise) in standard type IV rat cages (Ehret, Emmendingen, Germany) under a 12 h artificial light-dark cycle. Standard laboratory rat food (Ssniff, Soest, Germany) and tap water were provided ad libitum throughout the experimental period (unless stated otherwise). Female rats estrous cycle was not controlled for, as it has been reported that when rats were allowed to cycle freely no differences are seen in operant responding (Priddy et al., 2017; Roberts et al., 1998). All experimental procedures were approved by the Committee on Animal Care and Use (Regierungspräsidium Karlsruhe, Germany) and were carried out in accordance with the local Animal Welfare Act and the European Communities Council Directive of 22 September 2010 (2010/63/EU).

### Peer-rejection model

The study design was based on our previous model (Schneider et al., 2016a), where we examined the long-term consequences of peer-rejection-like ASEs in male and female rats on alcohol-seeking behaviors. In the present study Wistar rats were subjected to either control or peer-rejection condition throughout adolescence immediately after weaning (postnatal days, PD, 21-50). Control condition: male and female Wistar rats were housed in same sex groups of four under standard housing conditions in separate rooms. Peer-rejection condition: one Wistar rat was housed together with three age- and sex-matched Fischer 344 rearing partners. On PD 50, control Wistar rats were re-grouped with unfamiliar Wistar rats. While peer-rejected Wistar rats were grouped with unfamiliar Wistar rats from the same rearing condition, thereby terminating the peer-rejection procedure. After the peer-rejection period, animals were allowed to recuperate into adulthood after which all animals began alcohol self-administration training.

### Cue-induced reinstatement of alcohol-seeking

#### Operant alcohol self-administration apparatus

Cue-induced reinstatement of alcohol seeking was carried out in operant chambers (MED Associates Inc., St. Albans, VT) enclosed in ventilated sound-attenuating cubicles in control/peer-rejection female (n=20/20) and male (n=12/12) rats. The chambers were equipped with a response lever on each side panel of the chamber. Responses at the active lever activated a syringe pump that delivered a ∼30µl drop of fluid into a liquid receptacle next to it. Responses at the inactive lever were recorded but had no programmed consequences. A light stimulus (house light) was mounted above the left and right response levers of the self-administration chamber. An IBM compatible computer controlled the delivery of fluids, presentation of stimuli and data recording.

#### Alcohol self-administration acquisition and extinction phase

All animal training and testing sessions were performed during the dark phase of their light/dark cycle. Animals were trained to self-administer either 10% (v/v) alcohol or water in daily 30-min sessions using a fixed-ratio 1 (FR 1) schedule. The purpose of the acquisition phase was to train the animals to discriminate between the availability of alcohol (reinforcement) and water (non-reinforcement). Discriminative stimuli predicting alcohol or water availability were presented during each alcohol or water self-administration session (one 30-min session/day). An orange flavour extract served as the contextual stimulus (S+) for alcohol, whereas water availability was signalled by a lemon grass extract (S-). These olfactory stimuli were generated by depositing few drops of the respective extract into the bedding of the operant chamber before each session. In addition, each lever press resulting in alcohol delivery was accompanied by a 5-s blinking-light conditioned stimulus (CS+), whereas a 5-s constant-light stimulus (CS-) was presented with water delivery. The 5-s period served as a “time-out”, during which responses were recorded but not reinforced. At the end of each session, the bedding of the chamber was changed and trays were thoroughly cleaned. During the first two days of acquisition, animals were kept fluid deprived for 22 hours per day. Subsequently, alcohol and water sessions were conducted without fluid deprivation in a random manner until the animals received a total of 10 alcohol and 10 water sessions.

After completing the acquisition phase, rats were subjected to daily 30-min extinction sessions for five consecutive days, which in total was sufficient to reach reduced response rates approximating the extinction criterion of 20% of the last acquisition sessions. Extinction sessions began by extending the levers without presenting olfactory discriminative stimuli. Responses at the previously active lever activated the syringe pump, without resulting in the delivery of either alcohol or water or the presentation of response-contingent cues (stimulus blinking-light or constant-light).

#### Alcohol cue-induced reinstatement

Reinstatement testing began two days after the final extinction session. In these tests, rats were exposed to the same conditions as during the acquisition phase, except that the liquids (alcohol or water) were made unavailable. Sessions were initiated by the extension of both levers and the presentation of either the alcohol- (S+) or water- (S-) associated discriminative stimuli. Responses at the active lever were followed by the activation of the syringe pump and the presentation of the CS+ (blinking-light) in the S+ condition or the CS- (constant-light) in the S-condition. Half of animals were tested under the S+/CS+ condition on day 1 and under the S-/CS-condition on day 2. Conditions were reversed for the other half of animals. The number of responses on both the active (i.e., alcohol-associated lever for S+/CS+ condition and water-associated lever for S-/CS-condition) and inactive lever (i.e., water-associated lever for S+/CS+ condition and alcohol-associated lever for S-/CS-condition) was recorded throughout the experiment. Further behavioural testing took place for all animals after reinstatement testing.

### Circadian locomotor activity measurements by the E-motion system

In order to test home-cage locomotor activity control/peer-rejected female (n=12/12) and male (n=12/12) rats were separated into single-cages for three days and their number of movements was monitored by use of an infrared sensor connected to a recording and data storing system (Infra-e-motion, Henstedt-Ulzburg, Germany). Home cage locomotion was measured after alcohol reinstatement. An Infra-e-motion device was placed above each cage (30 cm from the bottom) so that the rat could be detected at any position inside the cage. The device was sampling every second whether the rat was moving or not. The sensor could detect body movement of the rat of at least 1.5 cm from one sample point to the next. The data measured by each Mouse-E-Motion device were downloaded onto a personal computer and processed with Microsoft Excel.

### Sucrose consumption test

To measure sucrose consumption control/peer-rejected female (n=12/12) and male (n=12/12) rats were kept single-housed. All rats were placed into separate single house cages. Two pre-weighed bottles, one containing tap water and the other containing the assigned sucrose solution, was placed on each cage. After 24 h, bottles were removed and re-weighed and liquid consumption was measured as the difference in the weight (g) of each bottle before and after the test as a parameter of hedonic responsiveness in rats (Willner et al., 1987). Furthermore, sucrose preference was measured as a parameter of hedonic state in rats, that was calculated using the formula: sucrose preference = [consumed sucrose solution/total liquid consumed (sucrose solution + water)] × 100. All rats were given ad libitum access to one bottle with tap water and another bottle with sucrose solution for 24 hours. Male rats received 0.35% (w/v) sucrose solution while female rats were given 0.15% (w/v) sucrose. Different concentration where used between sexes as female rats have been shown to be more sensitive to sweet tasting solutions (Curtis et al., 2004). The positions of bottles were counterbalanced to avoid location preferences and subsequent 24-hour intake of water and sucrose solution was measured.

### Statistical analysis

Data obtained from the self-administration and cue-induced alcohol-seeking experiments was analysed by use of a three-way ANOVA with repeated measures [between-subjects factors were: condition (control vs. peer-rejected) and within-subject factors: lever (active vs. inactive) and session (extinction vs. reinstatement)]. Locomotor activity data was analysed using 1-hour recordings using two-way ANOVA with repeated measures [factors were: group (control vs. peer-rejected) and time]. Furthermore, 24-hour locomotor activity, dark-phase and light-phase data was analyzed using a one-way ANOVA. One-way ANOVA was used for analysis of sucrose intake measures [factors were: group (control vs. peer-rejected) and solution (sucrose). Tukey’s honestly significant difference (HSD) post-hoc test was performed when appropriate. We used a two-sided significance level for all analyses and the assigned level of statistical significance was set to p < 0.05.

## Results

### Cue-induced reinstatement of alcohol seeking

Female control rats reached 97±13 alcohol-associated lever responses (S+/CS+ condition) and 21±2 water-associated lever responses (S-/CS-condition) while male control rats had 100±21 alcohol-associated lever responses (S+/CS+ condition) and 28±6 water-associated lever responses (S-/CS-condition) by the end of the acquisition phase. Female peer-rejected rats reached 124±12 alcohol-associated lever responses (S+/CS+ condition) and 29±4 water-associated lever responses (S-/CS-condition) and male peer-rejected rats had 80±16 alcohol-associated lever responses (S+/CS+ condition) and 16±2 water-associated lever responses (S-/CS-condition) by the end of the acquisition phase. Neither alcohol-nor water self-administration acquisition differed significantly between control and peer-rejected animals [water F(1,38) = 0.008, p = > 0.05 and Eth F(1,38) = 1.41, p > 0.05] in female rats and water [F(1,22) = 2.35, p >0.05] and Eth [F(1,22) = 0.86 p > 0.05] in male rats (Fig. 1).

**Figure 1.**
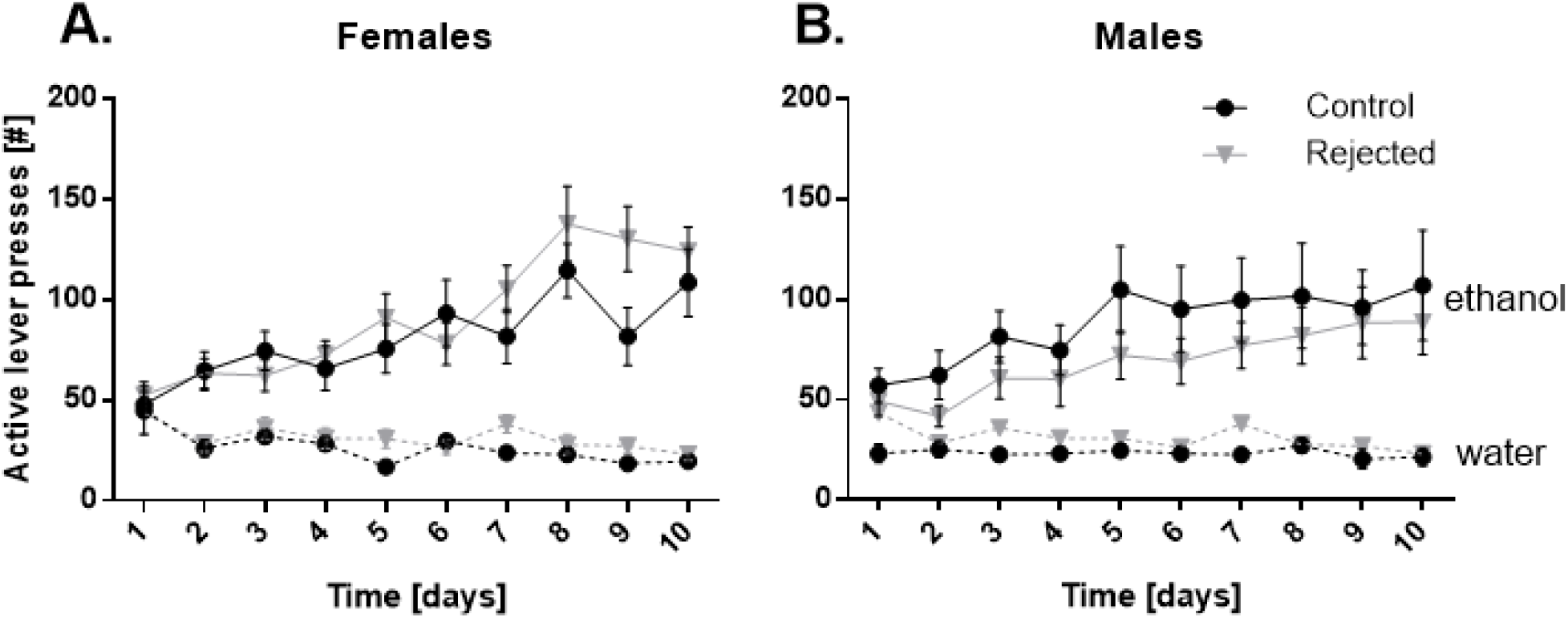
Average number of alcohol-paired (S+/CS+) and water-paired lever responses for each session (day) of self-administration under a FR1 schedule of reinforcement in females (n=20/20) (A) and males (n=12/12) (B). The alcohol-paired (ethanol) lever responses were greater than the (water) lever responses but no group differences were observed in alcohol or water consumption between controls (CTL) and peer-rejected (PR) animals. Data are presented as means ± S.E.M.

Three-way ANOVA with repeated measures revealed that all animals increased their pressing for the alcohol lever during the cue-induced reinstatement test compared to the last extinction session in female [F(1,38) = 110.60, p < 0.001] and male [F(1,22) = 41.0, p < 0.001] rats (Fig. 2). Peer-rejected female rats responded significantly more on the active lever during the alcohol cue-induced reinstatement test [F(1,38) = 7.35, p = < 0.01] (Fig. 2A). While an opposing effect was observed in males, where peer-rejected animals responded significantly less during the alcohol cue-induced reinstatement test [F(1,22) = 4.87, p < 0.05] (Fig. 2C). Tukey’s post-hoc analysis demonstrated that responding on the inactive lever was not significantly different between groups during the reinstatement test in either female or male animals (Fig. 2B, 2D respectively).

**Figure 2.**
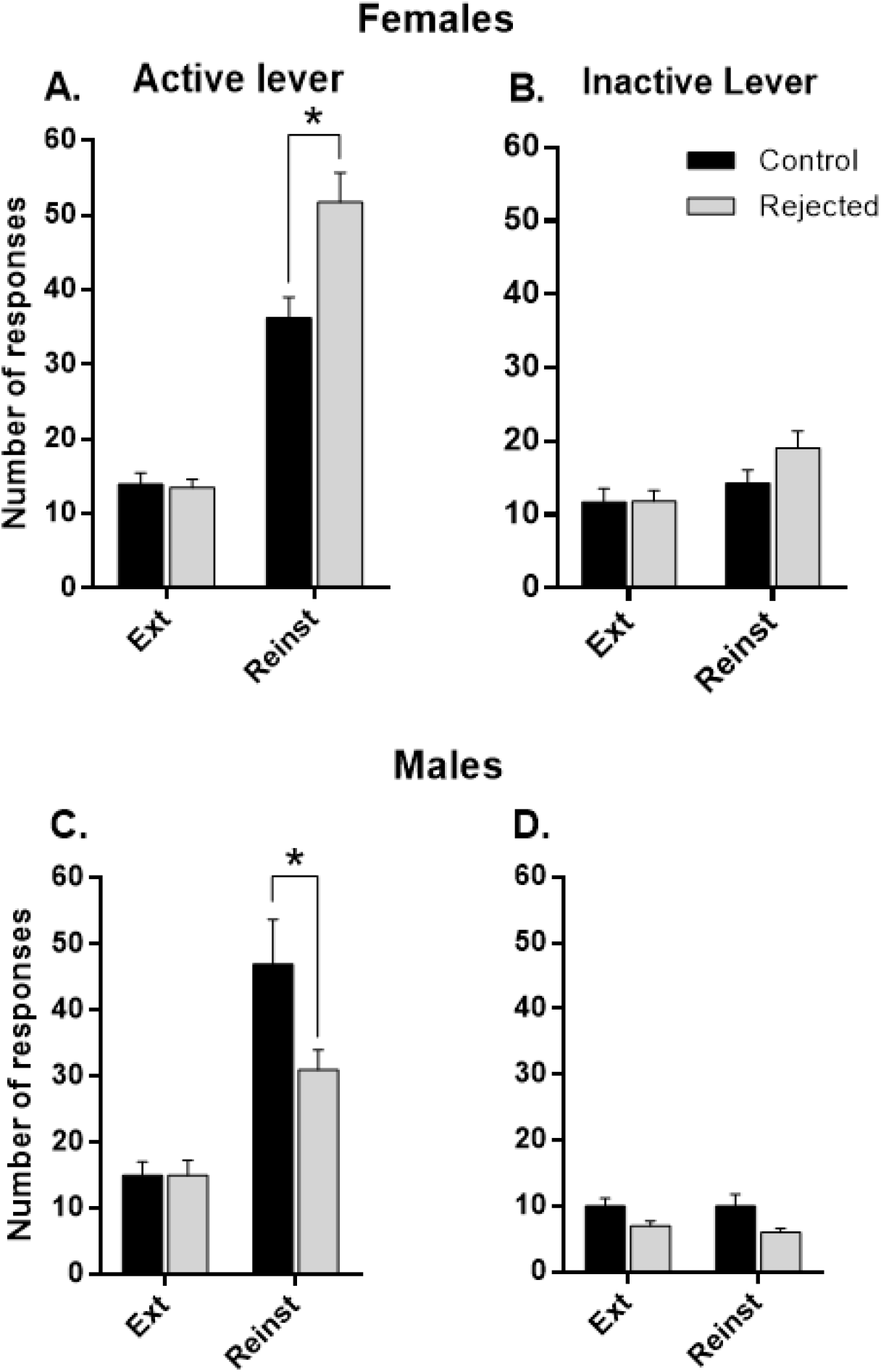
Alcohol cue-induced reinstatement in control and peer-rejected female (A-B, n=20/20) and male (C-D, n=12/12) rats. Data are shown as the average number of lever presses on the active (A, C) and inactive (B, D) levers during the last three extinction sessions (Ext) and as the number of responses after the presentation of stimuli previously paired with either alcohol (Reinst). Data are presented as means ± S.E.M. * indicates significant group differences, p <0.01.

Further post hoc analysis of the interaction effects revealed an effect of the housing condition (control vs. peer-rejected) on alcohol cue-induced reinstatement in both female [F(1,38) = 5.09, p < 0.05] and male [F(1,22) = 4.43, p < 0.05] rats. Responding during the water reinstatement test was also higher compared to the last extinction sessions in both female [F(1,38) = 142.54, p < 0.001] and male rats [F(1,22) = 6.09, p < 0.05] (data not shown).

### Home-cage circadian locomotor activity measurements by the E-motion system

A two-way ANOVA with repeated measures revealed that peer-rejection procedure had no significant effect on circadian locomotor activity across three days in female [F(1,22) = 0.009, p > 0.05] or male [F(1,22) = 0.78, p > 0.05] rats. A main effect of day was observed in male [F(1,22) = 17.84, p < 0.001] rats, however, no differences of day were observed in female [F(1,22) = 2.52, p > 0.05] rats. Tukey’s HSD revealed no significant between-group differences across days in male rats. Average 24-hour activity did not differ in either female [F(1,22) = 0.009, p > 0.92], or male [F(1,22) = 0.78, p > 0.05] rats (Fig. 3). All rats showed a rodent typical increase in locomotion during the dark phase and reduction of activity during the light phase. Analysis of dark phase (04:00–16:00) locomotor activity revealed no differences in female [F(1,22) = 0.38, p > 0.05] or male [F(1,22) = 0.42, p > 0.05] rats. Similarly, we found no circadian dysregulation in light phase (16:00–04:00) locomotor activity in either female [F(1,22) = 0.10, p > 0.05] or male [F(1,22) = 0.36, p > 0.05] rats.

**Figure 3.**
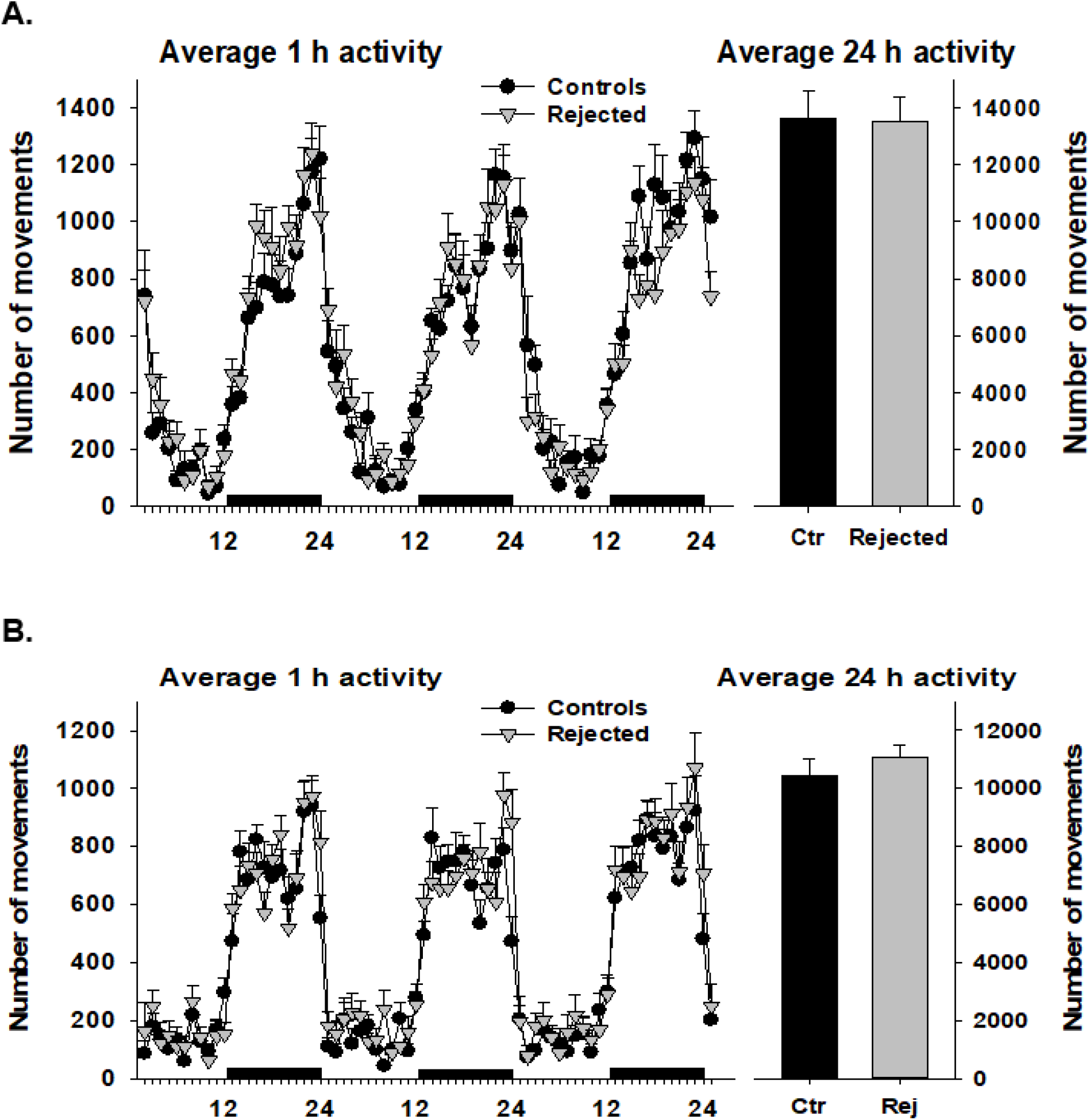
Circadian activity recordings in control and peer-rejected female (A) (n=12/12) and (B) (n=12/12) male rats measured as the number of movements during three consecutive days in the home-cage (black horizontal bars mark the dark (active) phases of the circadian cycle). Left figures demonstrate total number of movements during each consecutive hour and right figures represent total number of movements during 24-hours. All data are expressed as means ± SEM.

### Sucrose consumption test

Initial analysis of consumption of sucrose solution during the 24-hour free-choice sucrose preference test was significantly lower in rejected male rats compared to control animals [F(1,22) = 4.99, p < 0.05]. When sucrose consumption was corrected for weight these between group differences disappeared in males [F(1,22) = 3.23, p > 0.05] (Fig 4A). Total weight corrected sucrose consumption did not differ between female rejected and control rats [F(1,22) = 0.14, p > 0.05] (Fig 4B). No differences where observed in sucrose preference between groups in either sex.

**Figure 4.**
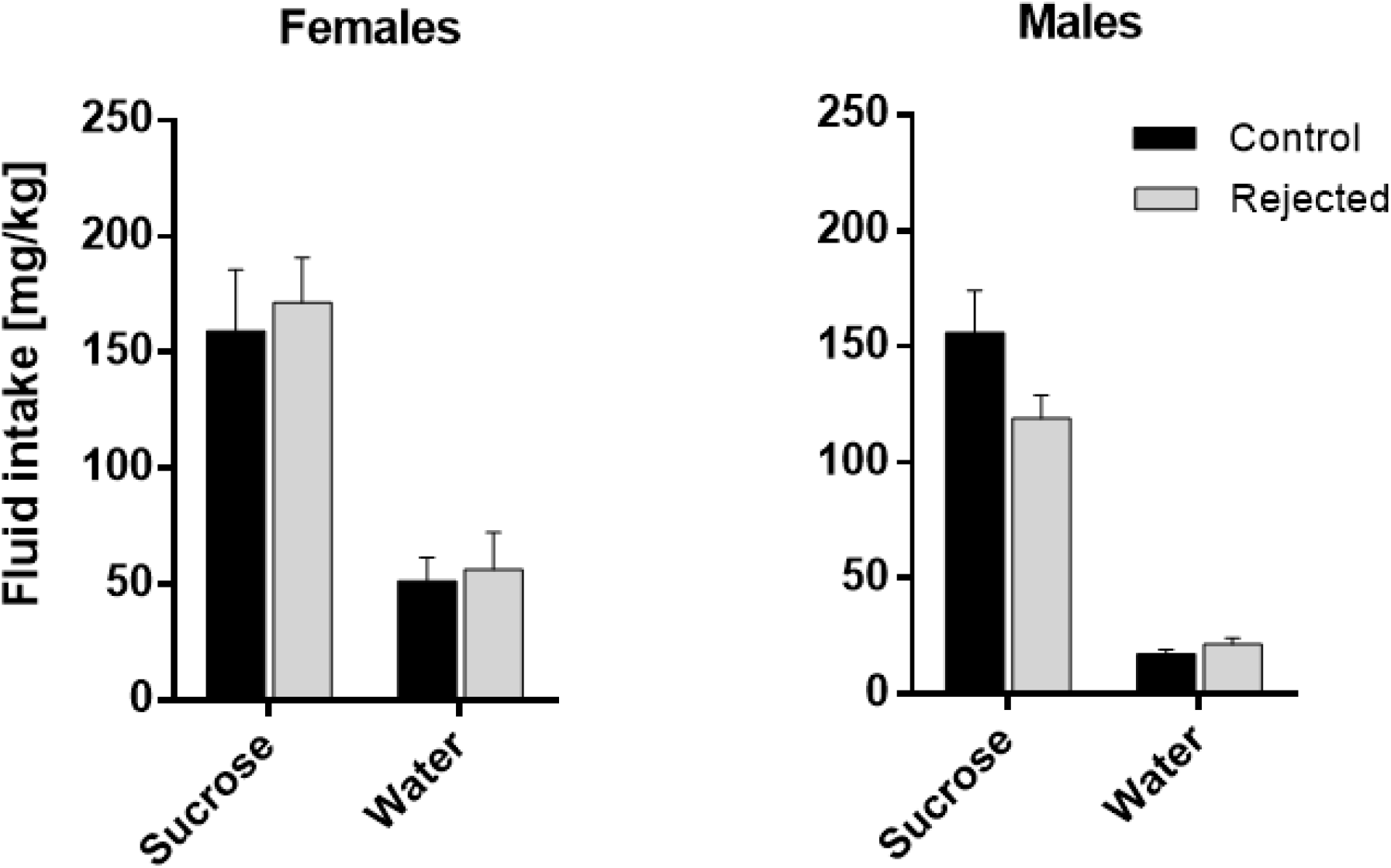
Intake of water and sucrose solution (mg/kg) in control and peer-rejected by (A) female (n=12/12) and (B) male (n=12/12) rats. All animal had free 24-hour access to a water bottle and a bottle containing either 0.15% sucrose solution (female rats) or 0.35% sucrose solution (male rats). All data are expressed as means ± SEM.

## Discussion

In this study, we investigated how peer-rejection-like ASEs across adolescence affected alcohol-seeking behavior, circadian locomotor activity and sucrose consumption in adult female and male Wistar rats. We observed no differences in either acquisition of alcohol and water self-administration or extinction learning between control and peer-rejected animal groups across sexes (Fig. 1). Interestingly, peer-rejection had sex-dependent effects on alcohol cue-induced reinstatement of alcohol-seeking. Specifically, our data showed that peer-rejection enhanced later life alcohol cue-induced reinstatement only in adult female Wistar rats. We observed the opposite effect of peer-rejection in males (Fig. 2). Our data are analogous to the human condition were women exposed to ACEs have been reported to have an increased risk for relapse (Heffner et al., 2011; Hyman et al., 2008; Kennedy et al., 2013). We conclude that the presented ecologically valid model of peer-rejection can be used in future studies to study the molecular and neurobiological underpinnings of persistent sex-dependent effects of adolescent ASEs on later alcohol-seeking and relapse-like behavior.

There is a gap in preclinical research on ASEs and their long-term impact on relapse-like behavior. We found a single study that has examined the long-term effects of ASEs on later-life alcohol relapse-like behavior in male and female rats (Bertholomey et al., 2016). Similar to our findings these authors demonstrated that pharmacological stress with corticosterone injections throughout adolescence increased cue-induced reinstatement more in females than males (Bertholomey et al., 2016). Interestingly, in our previous peer-rejection studies we observed alterations in corticosterone levels in adolescence (Schneider et al., 2014), together these data supports a role of corticosterone-mediated effects in later life alcohol-seeking behaviors. Corticosterone is a well-known modulator of stress as well as sleep (Hirotsu et al., 2015).

Emerging evidence has linked ACEs and inadequate social support to increased risk of circadian dysregulation (Kajeepeta et al., 2016; Kent et al., 2015). Circadian dysregulation in turn is associated with a myriad of health risks including increased alcohol consumption in both rodents and humans (reviewed by Jagannath et al., 2013). Moreover, alcohol can also influence circadian related gene expression and circadian rhythmicity (Perreau-Lenz and Spanagel, 2015). As our peer-rejection-like ASE model combines voluntary alcohol self-administration we assume it could impact circadian rhythms and induce circadian dysregulation. Our analysis initially revealed a main effect of day on locomotion, however post-hoc analysis demonstrated no significant differences between groups across days. This main effect of day was driven by a general increase in locomotion in both groups across subsequent days. Together these data indicate that peer-rejection does not induce long-term circadian dysregulation. Other studies that have reported circadian dysregulation after ASEs (Wells et al., 2017) have used more severe stressors such as chronic social defeat, this along with the fact that our circadian measurements were assessed 3 weeks after alcohol self-administration may explain the null findings in the present study.

Alongside circadian dysregulation, ACEs have been reported to induce alterations within the reward system. The cumulative stress of ACEs is often associated with a blunted reward response in humans (Hanson et al., 2016) and rodents (Willner et al., 1987) alike. Similar to individuals with ACEs, individuals with alcohol problems show similar neurobiological alterations in their reward processing (Novick et al., 2018) suggesting that some of the underlying neurobiological mechanisms of ACEs and AUD may overlap. For instance, both populations show a blunted response to reward in the ventral striatum, a region known to be involved in reward learning and decision-making (Cox and Witten, 2019; Wrase et al., 2007). We had previously demonstrated that peer-rejected rats have a decreased affinity for social reward (Schneider et al., 2016a), we were interested in assessing whether these deficits may be generalizable to palatable foods. Decreased preference for sweet tasting solutions is often used as a proxy for anhedonia-like behavior in rodent research (Papp et al., 1991). To assess for these deficits, we used the sucrose consumption test. We evaluated potential dysregulation of sweet reward sensitivity and anhedonia-like behavior by proxy of sucrose preference. However, we found no differences in sucrose consumption or sucrose preference in rejected female or male and control rats (Fig. 4). Notably, peer-rejected Wistar males exhibited a trend (p = 0.087) towards reduced sucrose consumption when compared to controls.

The observed changes in alcohol-seeking behavior reflects a quantitative sex-difference that may represent a vulnerability in rejected female rats and potentially a resilience in peer-rejected male rats to alcohol-seeking behaviors in adulthood. A potential driver of increased risk of alcohol reinstatement in females may be mediated by the neurobiological alterations reported by us previously (Schneider et al., 2016a, 2016b, 2014). Specifically, we found differences in the endocannabinoid system. We observed increased cannabinoid receptor 1(CB1R) activity in the amygdala and thalamus and decreased activity of fatty acid amide hydrolase (FAAH) in the amygdala. The endocannabinoid system and especially the CB1R is known to play an important part in regulating play and mood during adolescence (Schneider et al., 2015) and peer-rejection in turn seems to persistently alter this system potentially through altered play behaviors and via peer-rejection during adolescence (Schneider et al., 2016b). What remains unclear is how these alterations mediate enhanced alcohol-seeking behaviors in peer-rejected female rats. Numerous studies have demonstrated the importance of CB1R in alcohol-seeking behaviors. Where alcohol intake and alcohol-seeking behaviors are enhanced with CB1R agonists while on the other hand antagonists attenuate alcohol self-administration (Reviewed in: Fattore et al., 2007; Spanagel, 2020). Additionally, decreased FAAH levels have been observed in high alcohol preferring rats previously (Hansson et al., 2007). Together, these studies suggest that increased CB1R activity and decreased FAAH levels could mediate the observed enhancement in alcohol-seeking behaviors seen in rejected female rats.

In conclusion, female Wistar rats exposed to peer-rejection across adolescence show an increased propensity to relapse-like behavior similar to the human condition, while their peer-rejected male counterparts may demonstrate resilience-like effect to alcohol cue-induced reinstatement later in life. Although, our model cannot fully replicate the human condition of ACEs, it recapitulates behavioral deficits that impact alcohol-seeking behavior later in life. Finally, our model provides an approach with good face validity to studying the molecular and neurobiological mechanisms of ASEs on alcohol and potentially on other drugs of abuse across all phases of addiction-like behaviors in male and female rats.

## Acknowledgements/Funding

We would like to thank Sabrina Koch for the excellent technical assistance. This project was funded by the Deutsche Forschungsgemeinschaft (DFG, German Research Foundation, Project-ID 402170461-TRR26510 (Heinz et al., 2020) and Graduiertenkolleg GRK2350/1) and the Bundesministerium für Bildung und Forschung (BMBF, AERIAL program FKZ: 01EE1406C and SysMedSUDs FKZ: 01ZX1909).

